# Microbial contributions to subterranean methane sinks

**DOI:** 10.1101/034801

**Authors:** J. T. Lennon, D. Nguyễn-Thùy, N. Phạm Đú’c, A. Drobniak, P. Tạ Hòa, T.M. Pham, T. Streil, K.D. Webster, A. Schimmelmann

**Affiliations:** Department of Biology, Indiana University, Bloomington, Indiana, USA; Faculty of Geology, Vietnam National University, Hanoi, Vietnam; Department of Microbiology, Vietnam National University, Hanoi, Vietnam; Indiana Geological Survey, Indiana University, Bloomington, Indiana, USA; SARAD GmbH, Dresden, Germany; Department of Geological Sciences, Indiana University, Bloomington, Indiana, USA

## Abstract

Understanding the sources and sinks of methane (CH_4_) is critical for predicting and managing global biogeochemical cycles. Recent studies have reported that CH_4_ concentrations in cave ecosystems are depleted and that these subterranean environments may act as a daily sinks for atmospheric CH_4_. It has been hypothesized that this CH_4_ depletion may be caused by radiolysis, an abiotic process whereby CH_4_ is oxidized via interactions with ionizing radiation derived from radon decay. Alternatively, the depletion of CH_4_ concentrations could be due to biological processes, specifically oxidation by methanotrophic bacteria. We theoretically explored the radiolysis hypothesis and conclude that it is a kinetically constrained process that is unlikely to lead to the rapid loss of CH_4_ in subterranean environments. We present experimental results to support this claim. We tested the microbial oxidation hypothesis in a set of mesocosm experiments that were conducted in Vietnamese caves. Our results reveal that methanotrophic bacteria associated with cave rocks consume CH_4_ at a rate of 1.33 - 2.70 mg CH_4_ · m^-2^ · d^-1^. These CH_4_ oxidation rates equal or exceed what has been reported in other habitats, including agricultural systems, grasslands, deciduous forests, and Arctic tundra. As such, microbial methanotrophy has the potential to significantly oxidize CH_4_ in caves, but also smaller-size open subterranean spaces, such as cracks, fissures, and other pores that are connected to and rapidly exchange with the atmosphere. Future studies are needed to understand how subterranean CH_4_ oxidation scales up to affect regional and global CH_4_ cycling.

## Introduction

Atmospheric methane (CH_4_) is a potent greenhouse gas with rising concentrations that can mainly be attributed to anthropogenic activities (IPCC, 2013; US EPA, 2015). Credible forecasting of global warming by climate models mandates knowledge about the sources and sinks of atmospheric CH_4_. One potentially important, but overlooked sink of CH_4_ is the oxidation that occurs in subterranean environments. Recent studies have documented that cave ecosystems sometimes have subatmospheric concentrations of CH_4_. For example, in a four-year study of St. Michael’s Cave in Gibraltar, CH_4_ concentrations of cave air were typically 10-fold below atmospheric levels (Mattey *et al*., 2013). A similar pattern was documented in a set of Spanish caves with some samples having CH_4_ concentrations that were below detection limits suggesting near-complete removal of CH_4_ from underground air (Fernandez-Cortes *et al*., 2015).

Two hypotheses have been put forth to explain the pattern of CH_4_ depletion in subterranean environments. First, CH_4_ is a carbon and energy source that can be used by methanotrophic bacteria. Although methanotrophic bacteria were found in Movile Cave in Romania (Hutchens *et al*., 2004), microbiological surveys of methane oxidizing bacteria in caves are relatively uncommon. Inferences about methanotrophy in caves have also been made based on evidence from stable isotopes (Peryt *et al*., 2012). For example, an inverse relationship between CH_4_ concentrations and CH_4_ carbon stable isotope ratios (i.e., *δ*^13^C) in St. Michael’s cave was considered a diagnostic signature of methanotrophy (Mattey *et al*., 2013). A second hypothesis is that CH_4_ depletion in subterranean ecosystems is due to radiolysis. This abiotic mechanism of CH_4_ oxidation was developed to explain low CH_4_ concentrations in a poorly ventilated cave that had a high density of ions, but no recoverable methanotrophic bacteria (Fernandez-Cortes *et al*., 2015). An inverse correlation between the concentration of CH_4_ and ions in cave air suggests that α-particles from radon decay could contribute to removal of CH_4_ from subterranean environments (Fernandez-Cortes *et al*., 2015).

In this study, we evaluate the relative importance of biotic and abiotic mechanisms that have been put forward to explain low concentrations of CH4 observed in subterranean environments. First, we develop theoretical expectations in an effort to constrain the rates of radiolytic CH_4_ oxidation. Second, we present results from a controlled laboratory experiment aimed at quantifying the effect of ionizing radiation on the rate of CH4 oxidation. Third, we discuss findings from a set of field mesocosm experiments in Vietnamese caves to quantify the methanotrophic potential of cave microbial communities.

## Results and Discussion

**Weak theoretical support for the importance of radiolytic CH_4_ oxidation** — The following thought experiments reveal that radiolysis is a process that contributes minimally to CH_4_ oxidation in subterranean environments on the time scale of days to weeks. We arrive at this conclusion based on the imbalance between the large number of CH_4_ molecules and the comparatively small number of radioactive decay events that are typical in caves.

Ionizing radiation in the air of subterranean limestone-based ecosystems is derived predominantly from α-particles that are generated during radon decay (Cigna, 2005; Alvarez-Gallego *et al*., 2005). These α-particles could lead to the oxidation of CH_4_ *via* different mechanisms. For example, radiolysis could result from the direct collision of α-particles with CH_4_ molecules. In this case, an α-particle splits a CH_4_ molecule, which triggers a subsequent exothermic oxidation reaction of ions and radicals with atmospheric oxygen. However, with a decay rate of ˜35,000 ^222^Rn atoms per second in a cubic meter of air, as measured in a Spanish cave (Fernandez-Cortes *et al*., 2015), it would take nearly 50 million years to eliminate 2 ppm of CH_4_ as a result of direct collision between α-particles and CH_4_ molecules.

A more likely mechanism occurs when radiogenic energy interacts with water molecules and other major chemical constituents of cave air and thus produces ions and radicals that enter secondary chemical reactions with CH_4_. For example, radiolysis of water vapor *via* radon decay could generate hydroxyl radicals (•OH) that act to remove CH_4_. However, if every α-decay at 35,000 Bq m^-3^ generates 4.3 · 10^5^ ions and radicals (Fernandez-Cortes *et al*., 2015), it would still require more than 100 years to eliminate 2 ppm of CH_4_. In fact, this likely overestimates the potential for radiolytic CH_4_ oxidation since the calculations unrealistically assume that all •OH selectively react with CH_4_. In sum, our assumptions and calculations suggest that radiolysis is a kinetically constrained process that is unlikely to act as a daily CH_4_ sink in subterranean ecosystems (Fernandez-Cortes *et al*., 2015). More detail on the calculations that were used to arrive at our predictions can be found in the Supplementary Information,

**Weak experimental support for the importance of radiolytic CH_4_ oxidation** — Results from a laboratory experiment confirm our theoretical predictions by demonstrating that ionizing radiation had a minimal effect on CH_4_ oxidation rates. We placed 7.08 g uranium metal powder in a Petri dish on the bottom of a humid polyethylene bag containing 43 L of air with an elevated CH_4_ concentration (23.5 ppm). The radioactivity inside the closed bag containing depleted uranium was approximately 2.5 · 10^6^ Bq m^-3^, which is 70-fold higher than the natural radiation reported in Spanish cave air (Fernandez-Cortes *et al*., 2015). Yet, in the presence of strong ionizing radiation, CH_4_ was lost from the system at the slow rate of 0.197 ± 0.0005 (± standard error) ng CH_4_ · m^-3^ · d^-1^, which was indistinguishable from the diffusive loss of CH_4_ from polyethylene control bags lacking uranium (one-sample t-test: t_6_ =-0.97, *P* = 0.37, Fig. 1). More detail concerning experimental procedures can be found in Supplementary Information.

**Fig. 1.**
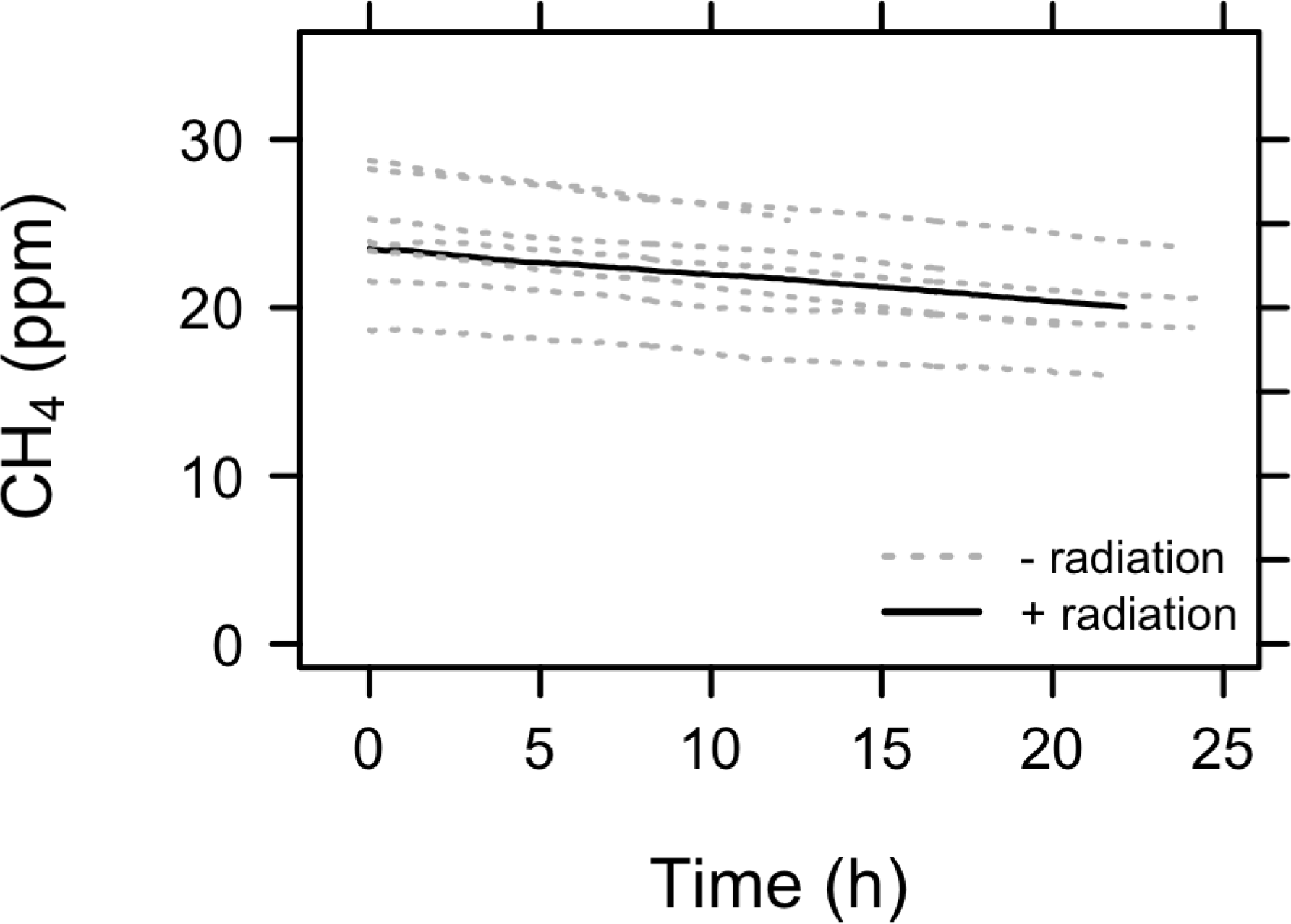
Rates of methane (CH_4_) oxidation were not significantly affected by ionizing radiation. We conducted a laboratory experiment where we tracked the concentration of CH_4_ in a polyethylene bag containing air and ionizing radiation from a source of uranium metal powder (black line, n = 1) to the concentration of CH_4_ in control bags without an added source of ionizing radiation (grey lines, n = 7). We attribute the slow loss of CH_4_ in all trials to diffusion through polyethylene bag.

**Strong experimental support for the importance of biotic CH_4_ oxidation** — The results from our field mesocosm experiments suggest that CH_4_ depletion can be achieved *via* the biological activity of methanotrophic bacteria that are associated with rocks inside cave ecosystems. In two separate caves with low radon abundances (≤100 Bq m^-3^) on the island of Cát Bà in Vietnam, we deployed 200-L polyethylene bags filled with cave air containing limestone rocks that were collected from inside the cave. Half of these mesocosms (n = 3) were treated with a 10 % bleach (sodium hypochlorite) solution to inhibit microbial activity ("dead") while the other mesocosms ("live") were treated with an equal volume of water (n = 3). After incubating *in situ* overnight, CH_4_ concentrations in the dead mesocosms were indistinguishable from the control mesocosms (no cave rocks) and the cave air (one-sample t-tests, *P* > 0.52, Fig. 2). In contrast, we observed an 87 % ± 0.047 % (mean ± SEM) reduction of CH_4_ concentrations in the live mesocosms.

**Fig. 2.**
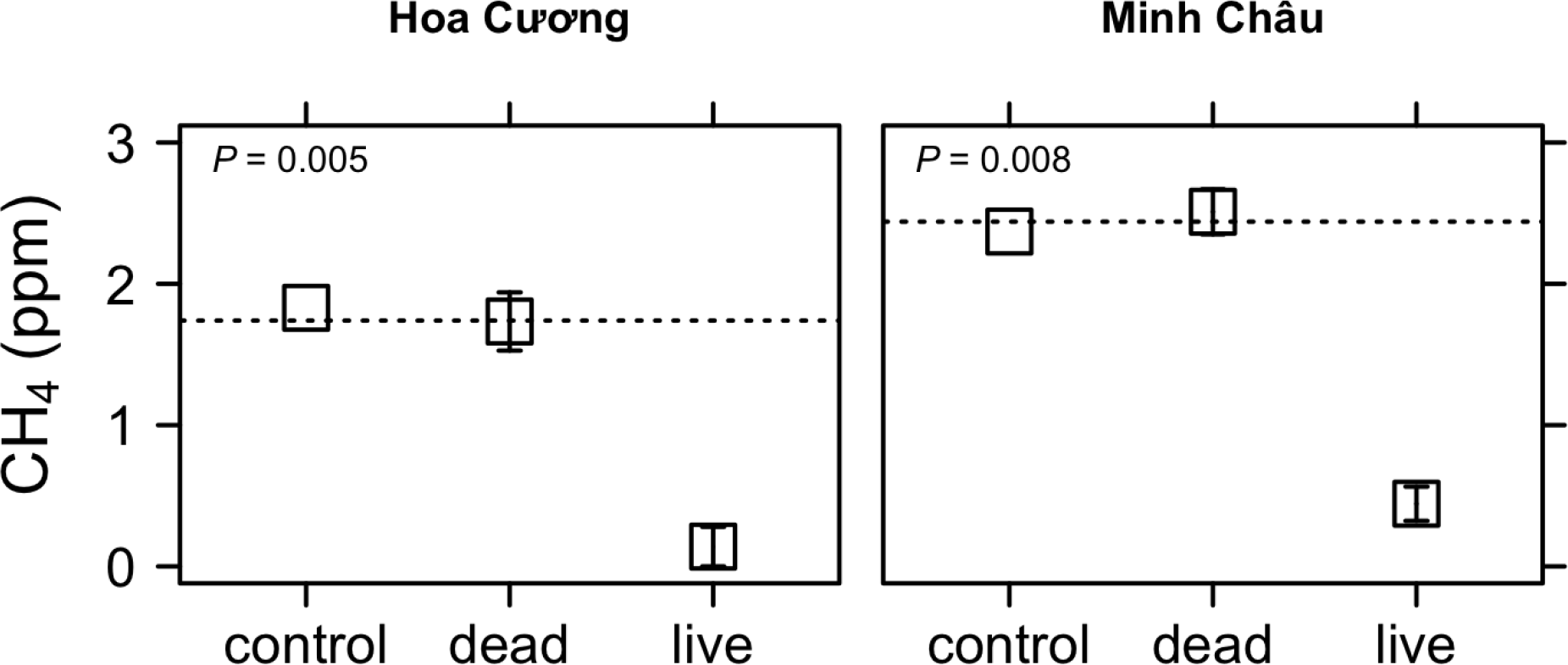
Field mesocosm experiments in two Vietnamese caves support the biological methane (CH_4_) oxidation hypothesis. Control mesocosms contained no cave rocks and provided an estimate for the diffusive loss of CH_4_; "dead" mesocosms contained cave rocks that were treated with a 10 % bleach solution; "live" mesocosms contained cave rocks and a volume of water (150 mL) equivalent to the volume of bleach used in the “dead” treatment. The dashed horizontal lines correspond to the CH_4_ concentrations in the Hoa Cuo’ng and Minh Châu caves on Cát Bà island, northern Vietnam.

From our experimental data, we estimate that the rate of CH_4_ oxidation associated with cave rocks was between 1.33 and 2.70 mg CH_4_ · m^-2^ · d^-1^. To the best of our knowledge, these are the first direct measurements of biological CH_4_ oxidation in a cave ecosystem. The magnitude of these rates equals or exceeds the rates of CH_4_ oxidation that have been reported in soils from agricultural systems, grasslands, mature forests, and Arctic tundra (Whalen & Reeburgh, 1990; Suwanwaree & Robertson, 2005; von Fischer *et al*., 2009). This comparison is noteworthy because caves maintain relatively constant temperatures thoughout the year, while soils in mid-to-high latitudes often experience lower temperatures during the winter season, which results in reduces rates of CH_4_ oxidation (e.g., Groffman *et al*., 2006). As such, when integrated over annual time scales, subterranean enviornments like caves may act as a larger sink for atmospheric CH_4_ than has been previously recognized.

Our experiments revealed that methanotrophic bacteria were abundant in the biofilms that were associated with Vietnamese cave rocks. We conducted quantitative PCR assays on DNA extracted from rocks that were incubated in the live mesocosms using primers that targeted the particulate methane monoxygenase (*pmo*A) gene, which is responsible for bacterial CH_4_ oxidation (see Supplementary Information for more detail). From this, we recovered 1.0 · 10^4^ to 1.5 · 10^4^ *pmo*A gene copies per gram of rock biofilm. When standardized by 16S rRNA gene copy number, we estimate that the relative abundance of methanotrophs in the cave biofilms ranged from 0.16 to 1.48 % of the microbial community.

Despite recent global-scale efforts to survey the diversity of microbial communities from a wide range of habitats, reports of methane oxidizing bacteria from cave ecosystems are scarce. For example, using cultivation-independent approaches, no sequences closely matching known methanotrophs were recovered from the Frasassi cave complex in central Italy (Macalady *et al*., 2006). Methanotrophs were recovered from some, but not all Spanish caves (Fernandez-Cortes *et al*., 2015). In limestone caves of Kartchner Caverns, Arizona (USA), a single sequence was recovered that was closely related to *Methylocella*, which is a facultative methanotroph (Ortiz *et al*., 2013). Similarly, only one sequence from the walls of a karstic cave in Slovenia was closely related to *Methylococcus*, which is an obligate methanotroph (Pašič *et al*., 2010). In contrast, the presence and activity of methanotrophs was documented in water and mat samples collected from Movile Cave using stable isotope probing (SIP). In this study, researchers tracked ^13^Clabeled CH_4_ into the DNA of bacteria that were closely related to known methanotrophs such as *Methylomonas, Methylococcus*, and *Methylocystis/Methylosinus* (Hutchens *et al*., 2004). Given their potential role in consuming subterranean CH_4_, more studies are needed to characterize the diversity and activity of methanotrophs in a wider range of cave ecosystems.

In the methane-depleted Castañar Cave in Spain, the importance of methanotrophy was ruled out based on the assumption that bacteria would not be able to meet their metabolic demands for maintenance and growth (Fernandez-Cortes *et al*., 2015). However, this argument overlooks two important ecophysiological features of microorganisms in natural ecosystems. First, growing evidence suggests that many microorganisms can tolerate extreme energy limitation on timescales ranging from centuries to millennia (Hoehler & Jørgensen, 2013) owing to life-history strategies such as dormancy (Lennon & Jones, 2011). Second, microorganisms in nature are commonly challenged with "feast or famine" conditions. For example, the supply of CH_4_ to cave habitats varies through time depending on the source of CH_4_, seasonality, ventilation, microclimatic conditions, and geography. Methanotrophic bacteria in caves are likely adapted to such fluctuations in CH_4_ concentrations, which are not captured with synoptic sampling.

**Conclusion** — Although ionizing radiation can accumulate in poorly vented, deep recesses of some caves, this is neither necessary nor sufficient to explain the pattern of CH_4_ depletion in subterranean ecosystems. Both theoretical and experimental lines of evidence suggest it is unlikely that radiolytically induced CH_4_ oxidation serves as a significant mechanism for rapid depletion of CH_4_ in cave air. Rather, our results support the hypothesis that bacterial methanotrophy alone has the potential to significantly oxidize CH_4_ in caves, but also smaller-size open subterranean spaces, such as cracks, fissures, and other pores that are connected to the atmosphere.

## Acknowledgements

We acknowledge support from the U.S. Department of Energy, Office of Science, Office of Basic Energy Sciences (DE-SC0006978 to A.S.); the National Science Foundation (1442246 to J.T.L); the U.S. Army Research Office (W911NF-14-1-0411 to J.T.L.); the Indiana University Office for the Vice President of International Affairs; the Indiana University College of Arts and Sciences; and the Indiana University Provost’s Travel Award for Women in Science. We thank G. Etiope for constructive feedback on an earlier version of this manuscript and G. Crouch for discussions about radiation physics. In addition, we thank B. K. Lehmkuhl for technical support, along with Minh Schimmelmann and Bui Thi Viet Ha for logistical support. Corresponding data and code for this manuscript can be found at https://github.com/LennonLab/radiolyticCH4.

## References

Alvarez-Gallego M, Garcia-Anton E, Fernandez-Cortes A, Cuezva S, Sanchez-Moral S. High radon levels in subterranean environments: monitoring and technical criteria to ensure human safety (case of Castañar cave, Spain) (2005). Journal of Environmental Radioactivity 145, 19–29.

Cigna AA (2005) Radon in caves. International Journal of Speleology 34, 1–18.

Fernandez-Cortes A, Cuezva S, Alvarez-Gallego M, Garcia-Anton E, Concepcion P, Benavente D, Jurado V, Saiz-Jimenez C, Sanchez-Moral S (2015) Subterranean atmospheres may act as daily methane sinks. Nature Communications 6, 7003.

Groffman PM, Hardy JP, Driscoll CT, Fahey TJ (2006) Snow depth, soil freezing, and fluxes of carbon dioxide, nitrous oxide and methane in a northern hardwood forest. Global Change Biology 12, 1748–1760

Hoehler TM, Jørgensen BB (2013) Microbial life under extreme energy limitation. Nature Reviews Microbiology 11, 83–94.

Hutchens E, Radajewski S, Dumont MG, McDonald IR, Murrell CJ (2004) Analysis of methanotrophic bacteria in Movile Cave by stable isotope probing. Environmental Microbiology 6, 111–120.

IPCC (2013) The Physical Science Basis. Contribution of Working Group I to the Fifth Assessment Report of the Intergovernmental Panel on Climate Change (eds. Stockner TF et al.) Cambridge University Press.

Lennon JT, Jones SE (2011) Microbial seed banks: ecological and evolutionary implications of dormancy. Nature Reviews Microbiology 9, 119–130.

Macalady JL, Lyon EH, Koffman B, Albertson LK, Meyer K, Galdenzi S, Mariani S (2006) Dominant microbial populations in limestone-corroding stream biofilms, Frasassi cave system, Italy. Applied and Enviornmental Microbiology 72 5596–5609.

Mattey DP, Fisher R, Atkinson TC, Latin J-P, Durrel R, Ainsworth M, Lowry D, Fairchild IJ (2013) Methane underground air in Gibraltar karst. Earth and Planetary Science Letters 374, 71–80.

Ortiz M, Neilson JW, Nelson WM, Legatzki A, Byrne A, Yu Y, Wing RA, Soderulund CA, Pryor BM, Pierson LS, Maier RM (2013) Profiling bacterial diversity and taxonomic composition on speleothem surfaces in Kartchner Caverns, AZ. Microbial Ecology 65 371–383.

Pašič L, Kovče B, Sket B, Herzog-Velikonja B (2010) Diversity of microbial communities colonizing the walls of a Karstic cave in Slovenia. FEMS Microbiology Ecology 71 50–60

Peryt TM, Kurakiewicz T, Peryt D, Poberezhskyy A (2012) Carbon and oxygen isotopic composition of the Middle Miocene Badenian gypsum-associated limestones of West Ukraine. Geologica Acta 10, 319–332.

Suwanwaree P, Robertson GP (2005) Methane oxidation in forest, successional, and no-till agricultural ecosystems: effects of nitrogen and soil disturbance. Soil Science Society of America Journal 69, 1722–1729

US EPA (US Environmental Protection Agency) (2015) Inventory of U.S. greenhouse gas emissions and sinks: 1990–2013.

von Fischer JC, Butters G, Duchateau PC, Thelwell RJ, Siller R (2009) In situ measures of methanotroph activity in upland soils: A reaction-diffusion model and field observation of water stress. Journal of Geophysical Research 114, 0148-0227/09/2008JG000731

Whalen SC, Reeburgh WS (1990) Consumption of atmospheric methane by tundra soils. Nature 346, 160–162.

